# Evidence that metastases arise *de novo* as new cancers from cells of target organs

**DOI:** 10.1101/2023.08.09.552586

**Authors:** Gorantla V. Raghuram, Kavita Pal, Soniya Shende, Naveen Kumar Khare, Vishalkumar Jadhav, Sushma Shinde, Ratnam Prasad, Harshada Wani, Indraneel Mittra

**Affiliations:** Translational Research Laboratory, Tata Memorial Centre, Advanced Centre for Treatment Research and Education in Cancer, Kharghar, Navi Mumbai – 410210, India; Homi Bhabha National Institute, Anushakti Nagar, Mumbai - 400 094, India

**Keywords:** circulating tumour cells, cell death, cell-free chromatin particles, hallmarks of cancer, immune check-points, blood-brain barrier

## Abstract

**Introduction:** Based on our earlier finding that cell-free chromatin particles (cfChPs) released from dying cancer cells are potentially oncogenic, we hypothesised that metastases arise as new cancers from the cells of target organs transformed by cfChPs released from dying cancer cells.

**Methods:** We fluorescently dually-labelled MDA-MB-231 human breast cancer cells and A-375 human melanoma cells in their DNA with BrdU and in their histones with CellLight® Histone 2BGFP. One hundred thousand dually-labelled cells were intravenously injected into SCID mice. In other experiments unlabelled cells were injected for detection of lung metastasis. Also intravenously injected were 700 ng of purified cfChPs isolated from radiation-killed MDA-MB-231 cells.

**Results:** We observed that the fluorescently dually-labelled MDA-MB-231 and A-375 cells died upon reaching the lungs and released dually-labelled fluorescent chromatin particles that accumulated in the nuclei of lung cells at 48 h. Injection of unlabelled cfChPs led to the activation of 10 hallmarks of cancer and immune checkpoints in lung cells at 72 h, suggesting that the lung cells had rapidly transformed into incipient cancer cells. Fluorescent *in situ* hybridisation analysis of lung metastases that subsequently developed using mouse and human specific DNA probes revealed that the tumour cells contained both mouse and human DNA in almost equal proportions. Similarly, immune-fluorescence analysis using species specific mAbs revealed that the tumour cells co-expressed mouse and human specific proteins. Metaphase preparations and single-cell clones developed from cell cultures of lung metastases were found to contain chimeric chromosomes containing both mouse and human DNA, and the cells to co-express both human- and mouse-specific proteins. The Intravenously injected purified cfChPs isolated from radiation-killed MDA-MB-231 cells also induced lung metastasis which predominantly contained mouse DNA strongly suggesting that the metastatic tumours had arisen from the mouse lung cells.

**Conclusion:** These results provide strong evidence that cfChPs released from dying cancer cells integrate into genomes of target lung cells to transform them into new cancers that masquerade as metastasis. They support our hypothesis that metastases arise from the cells of target organs and not from those of the primary tumour. These findings have implications for the principles of cancer therapy.

## Introduction

Metastasis is a complex process and the leading cause of cancer-related deaths. Despite extensive research over many decades, the mechanisms underlying the metastatic cascade are not well understood. The multiple processes involved in the metastatic cascade have been extensively reviewed **1,2)** and comprise of the following steps: 1) detachment of cells from the primary tumour and their intravasation into the circulatory / lymphatic system; 2) the intrvasated cells surviving immune attack by macrophages, natural killer cells and T cells; 3) surviving the shearing force of circulation; 4) navigating the vascular capillaries that are often narrower than the diameter of the tumour cells themselves; 5) exiting the vascular system to enter into the neighbouring normal tissues; 6) in case of the brain, having to cross the blood-brain barrier to extravasate into the brain parenchyma; 7) survive in an inhospitable environment, proliferate and colonize to form macroscopic metastases. Many researchers have investigated the complex molecular mechanisms underlying each of the above steps of the metastatic cascade **(see refs. 3 and 4 for review).** Yet metastasis continues to remain a poorly understood phenomenon and a major cause of death from cancer. Herein we present results that support a radically different model and suggest novel therapeutic possibilities.

The symbiotic influence of two deeply entrenched theories has dominated metastasis research for decades. The first theory suggests that metastatic tumours are histologically similar to the underlying primary tumours. Although this theory has not been scientifically tested, it continues to form the conceptual foundation of metastatic research. A recent study that evaluated this theory in a blinded experiment raised doubts about its veracity **(5)**. Nonetheless, this entrenched theory prompted the formulation of a second theory, which posits that cells leaving the primary tumour enter the circulation and travel to distant organs where they colonise to form metastases that are histologically similar to the primary tumour. The striking finding of Fidler **(6)** that 99% of cancer cells injected intravenously into mice die by the time they reach distant target organs was construed to suggest that the remaining 1% surviving cells are responsible for inducing metastasis. This conjecture led to the emergence of the theory of “metastatic inefficiency” **7,8)**, which ensured that the two entrenched theories remained unchallenged. The possibility that the 99% of the cells that die may have consequences for the development of metastasis remains unaddressed.

In accordance with Fidler’s report, we had earlier observed that intravenously injected cancer cells die upon reaching vital organs **(9)**. The fragmented chromosomes in the form of cell-free chromatin particles (cfChPs) that emerge from the dying cells accumulate in the nuclei of vital organs (lung, liver, brain and heart). Nuclear accumulation of cfChPs leads to induction of two critical hallmarks of cancer viz. DNA damage (genomic instability) and inflammation **(9)**. As simultaneous DNA damage and inflammation is a potent stimulus for oncogenic transformation, we had hypothesized that cfChPs from dying cancer cells can potentially transform cells of distant organs providing an alternative mechanism of cancer metastasis **(9)**. A similar model has also been proposed by several other investigators who have implicated horizontal transfer of nucleic acids released from dying cancer cells in the metastatic process **(10-14)**, leading to the hypothesis of “geno-metastasis” **11,12)**.

In the present study, we show that intravenously injected human cancer cells into severe combined immune-deficient (SCID) mice release cfChPs that accumulate in the nuclei of lung cells. This leads to the activation of 10 hallmarks of cancer and the immune checkpoint PD-L1 as early as 72 h following injection, suggesting that the lung cells are rapidly transformed into incipient cancer cells. The metastasis that subsequently developed contained both mouse and human specific DNA and the tumour cells were comprised of chimeric chromosomes containing DNA of both species. These findings indicated that the cfChPs of human origin had integrated into the genomes of mouse lung cells and transform them into new cancers that appeared as metastases. Evidence that cfChPs were indeed the agents responsible for transforming lung cells was strongly suggested by the demonstration that the intravenous injection of purified cfChPs isolated from radiation-killed human cancer cells also generated lung metastases which contained both mouse and human DNA.

## Methods

### Ethics approval

The experimental protocol was approved by the Institutional Animal Ethics Committee (IAEC) of the Advanced Centre for Treatment, Research, and Education in Cancer (ACTREC), Tata Memorial Centre (TMC), Navi Mumbai, India. The experiments were conducted in compliance with the ethical regulations and humane endpoint criteria of the IAEC and ARRIVE guidelines.

### Mice

Inbred female non-obese diabetic (NOD) mice (NOD.Cg-Prkdcscid/J) with severe combined immunodeficiency (SCID) were used in this study. They were obtained from the Institutional Animal Facility, maintained according to IAEC standards, and housed in pathogen-free cages containing husk bedding under a 12-h light/dark cycle with free access to water and food. A HVAC system was used to control room temperature, humidity, and air pressure. The ACTREC-IAEC maintains respectful treatment, care, and use of animals in scientific research and aims to contribute to the advancement of knowledge following ethical and scientific necessity. All scientists and technicians involved in this study underwent training in the ethical handling and management of animals under the supervision of Federation of European Laboratory Animal Science Associations (FELASA)-certified attending veterinarians. The animals were killed at appropriate time points by cervical dislocation under CO_2_ anaesthesia under the supervision of FELASA-trained animal facility personnel.

### Cell culture

We used two human cancer cell lines: MDA-MB-231 (human breast cancer) and A-375 (human malignant melanoma). Cells were obtained from the American Type Culture Collection (ATCC) and grown in Dulbecco’s modified Eagle medium (DMEM) with 10% fetal bovine serum (FBS) + 1% antibiotic cocktail in an atmosphere of 5% CO_2_ and 100% humidity.

### Fluorescent dual-labelling of cells

The MDA-MB-231 and A-375 cells were fluorescently dually-labelled according to a protocol previously described by us **(15)**. Cells were plated at a density of 6 × 10^4^ cells in Dulbecco’s Modified Eagle Medium, and after overnight culture (cell count 1 × 10^5^), their DNA was labelled using BrdU (10 μM / 1.5 ml for 24 h). BrdU was obtained from Sigma Chemicals, St. Louis, MO, USA; Catalogue No. B5002; Histones were labelled using CellLight^®^ Histone 2BGFP (60 µL / 1.5 ml for 36 h). CellLight^®^ Histone 2BGFP was obtained from Thermo Fisher Scientific, Waltham MA, USA; Catalogue No. C10594. Representative images of the fluorescently double-labelled MDA-MB-231 and A-375 cells are shown in Figure 1a.

**Figure 1:**
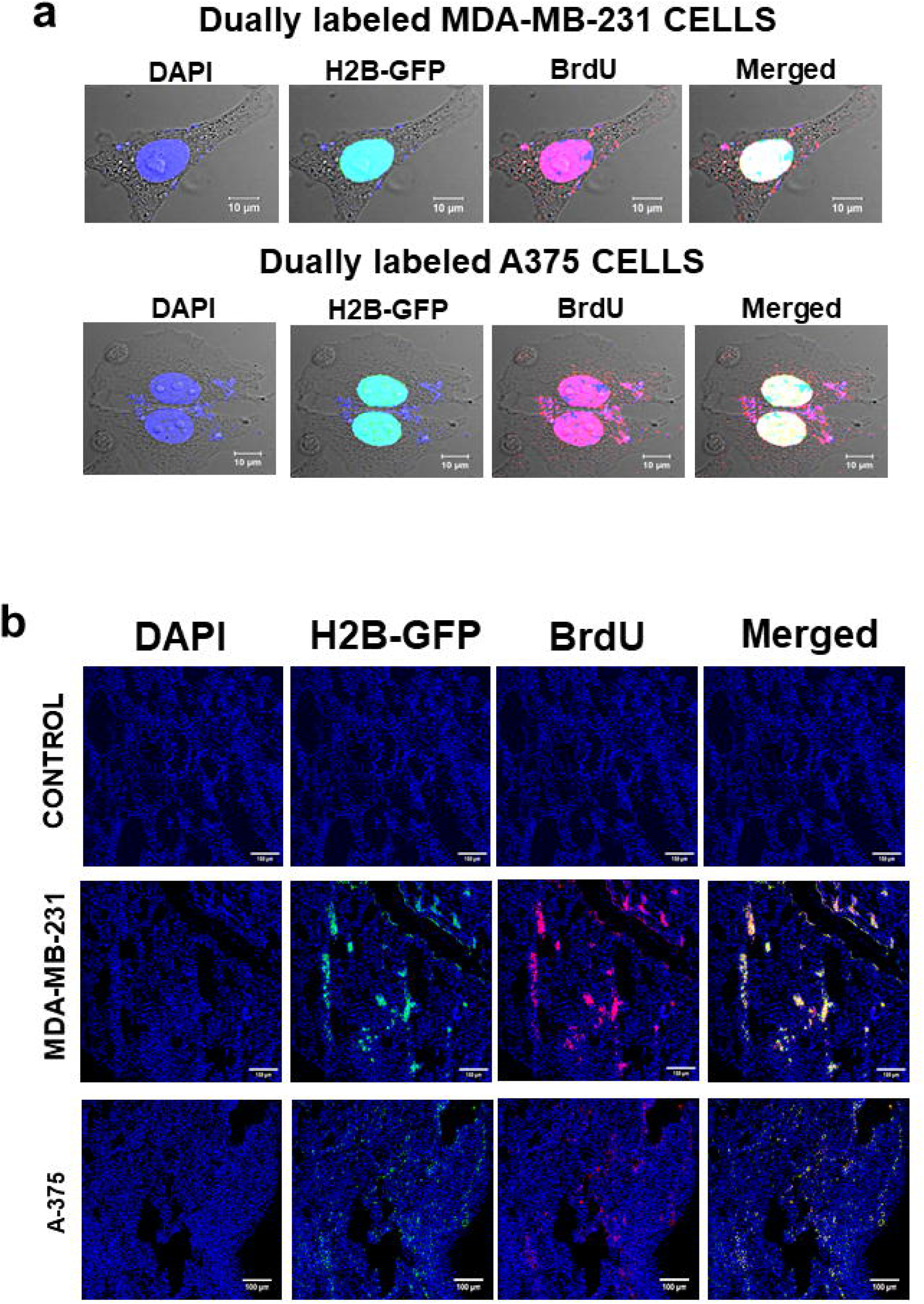
Representative images of fluorescently dual-labelled MDA-MB-231 and A-375 cells, and the accumulation of dual-labelled cfChPs in the lung cells of mice following intravenous injection; **a.** Representative confocal images of fluorescently dual-labelled MDA-MB-231 and A-375 cells, in which DNA was labelled with BrdU (red) and histones were labelled with H2B-GFP (green). The extra-nuclear DAPI and BrdU signals represent mitochondrial DNA. **b.** Fluorescence microscopy images showing patchy accumulation of dual-labelled particles representing cfChPs in lung cells following intravenous injections of fluorescently dually-labelled MDA-MB-231 and A-375 cells at 48 h. One hundred thousand cells were injected intravenously into SCID mice in both the cases.

### Intravenous injection and detection of fluorescently dual-labelled particles in lung cells

Dually labelled MDA-MB-231 and A-375 cells (1 × 10^5^) were intravenously injected into SCID mice, and the animals were sacrificed after 48 h. The lungs were harvested and cryosections were prepared for immunofluorescence (IF) staining with an anti-BrdU antibody (Abcam, Cambridge, UK; Catalogue no. ab70328). Images were acquired and analysed using an Applied Spectral Bioimaging System (ASI, Israel).

### Detection of hallmarks of cancer in lung cells

One hundred thousand unlabelled MDA-MB-231 cells were intravenously injected into SCID mice, while control SCID mice were injected with saline. Mice in both groups were sacrificed at 72 h, the lungs were harvested, and cryosections were prepared for IF analysis to detect the hallmarks of cancer, as defined by Hanahan and Weinberg **(16)**. Antibodies against 13 biomarkers representing 10 hallmarks were used in the analysis. Details of the antibodies used are provided in Supplementary Table 1 B and 1C. IF analysis was performed as previously described by us **(17)**. Images were acquired and analysed using an Applied Spectral Bioimaging System (ASI, Israel).

### Detection of lung metastasis

The MDA-MB-231 and A-375 cells induce lung metastasis when injected intravenously into SCID mice **18,19)**. Ten SCID mice were intravenously injected with 1 × 10^5^ MDA-MB-231 cells, and five mice were injected with an equal number of A-375 cells. Two mice from the MDA-MB-231 group and one mouse from the A-375 group were euthanised on days 37, 51, 76, 112, and 125. Formalin-fixed paraffin-embedded (FFPE) sections of the lungs were prepared, stained with haematoxylin and eosin (H&E), and examined for the histological detection of metastases. In the MDA-MB-231 group, grade III metastases were detected on day 112 in two mice and on day 125 in two mice (total number of metastases 4/10). One of the tumours detected on day 112 was used for fluorescence in situ hybridisation (FISH) and immuno-fluorescence (IF) analysis, and the other was used for preparing primary cultures. In the A-375 group, grade III metastases were detected in one mouse each on days 76, 112, and 125 (number of metastases 3/5). The tumour detected on day 76 was used for FISH and IF analysis and that detected on day 112 was used to prepare primary cultures. Representative H&E images of MDA-MB-231 and A-375 lung metastases are shown in Supplementary Figure 1.

### Development of primary cultures and single-cell clones

Lung metastases which were visible as white patches on the lung surface were carefully excised under aseptic conditions, minced with scalpel followed by collagenase I treatment (1mg/ml) for 30 min at 37^0^C with gentle shaking. The dissociated cells were centrifuged, washed X 2 with PBS and plated in 65mm^3^ petri dishes containing DMEM with 20% fetal bovine serum (FBS) + 1% antibiotic cocktail. The medium was changed after 24hr and cells were allowed to grow to confluency and passaged several times. After the 3-4 passages foci of the transformed cells became visible. Cells at the foci were carefully picked, trypsinised, and subcultured in the above medium. Single-cell clones were prepared from primary cultures using the serial dilution method and were used for further analysis. The epithelial nature of the primary cultures and single-cell clones were analysed using a recombinant anti-EpCAM antibody (EPR20533-63; Catalogue number: ab221552, Abcam; Supplementary Figure 2).

### Detection of species-specific DNA in lung metastasis by FISH

Unstained FFPE sections of metastatic tumours were processed for FISH analysis using mouse (green) or human (red)-specific whole genomic DNA probes (custom synthesised by ASI, Israel) as described by us previously **(17)** (Supplementary Table 1A). Images were acquired and analysed using an Applied Spectral Bioimaging System (ASI, Israel). At the outset, we performed extensive control experiments to ensure that the human and mouse FISH probes did not cross-react (Supplementary Figure 3).

### Detection of species-specific proteins in lung metastasis by IF

Unstained FFPE sections of metastatic tumours were processed for IF analysis using mouse- and human-specific antibodies (green and red, respectively) and appropriate secondary antibodies. We used species-specific monoclonal antibodies (mAbs), namely, HLA class 1-ABC, which is unique to humans, and MHC class II, which is unique to mice. Initially, we performed elaborate control experiments to ensure that the human and mouse mAbs did not cross-react (Supplementary Figure 3). The details of the antibodies used are provided in Supplementary Table 1B and 1C.

### Isolation of cfChPs from irradiated dying cancer cells

The MDA-MB-231 and A-375 cells grown in DMEM containing 10% foetal bovine serum were plated at a density of 1.5 × 10^6^ in 10 cm culture dishes. The following day (cell density ∼3 × 10^6^), cells were irradiated with γ rays (10 Gy) and incubated for 72 h. The floating dead cells in the supernatant were collected by centrifugation and used for cfChP isolation as described in detail by us previously **(17)**. An electron microscopy image of cfChPs isolated from the MDA-MB-231 cells is shown in Supplementary Figure 4.

### Intravenous injection of purified cfChPs into SCID mice

Five SCID mice were intravenously injected with 700 ng cfChPs isolated from dying MDA-MB-231 cells, and four mice were intravenously injected with cfChPs isolated from dying A-375 cells (2/4 mice in the A-375 group subsequently died). We chose to inject 700 ng of isolated cfChPs as this is the amount of DNA present in 100,000 injected cells. The animals were sacrificed after 150 days, and H&E sections of their lungs were examined for the development of metastases. Two out of five mice from the MDA-MB-231 group and 0 / 2 from the A-375 group harboured lung metastases. A representative H&E image of lung metastasis that developed following the intravenous injection of cfChPs isolated from MDA-MB-231 cells is shown in Figure 6a. FFPE sections were prepared and processed for FISH and IF as described above.

### Statistical analysis

Statistical analysis pertains to the 13 cancer hallmark biomarkers mentioned above. The mean (± SEM) values of biomarkers in the control and MDA-MB-231-injected groups were individually compared using two tailed Student’s t test. A combined analysis of the median values of all 13 biomarkers in control and MDA-MB-231 injected groups was performed using the Mann–Whitney U test (see the legend to Figure 2). Analyses were performed using GraphPad Prism Version 6.0. The threshold for statistical significance was set at p<0.05.

**Figure 2:**
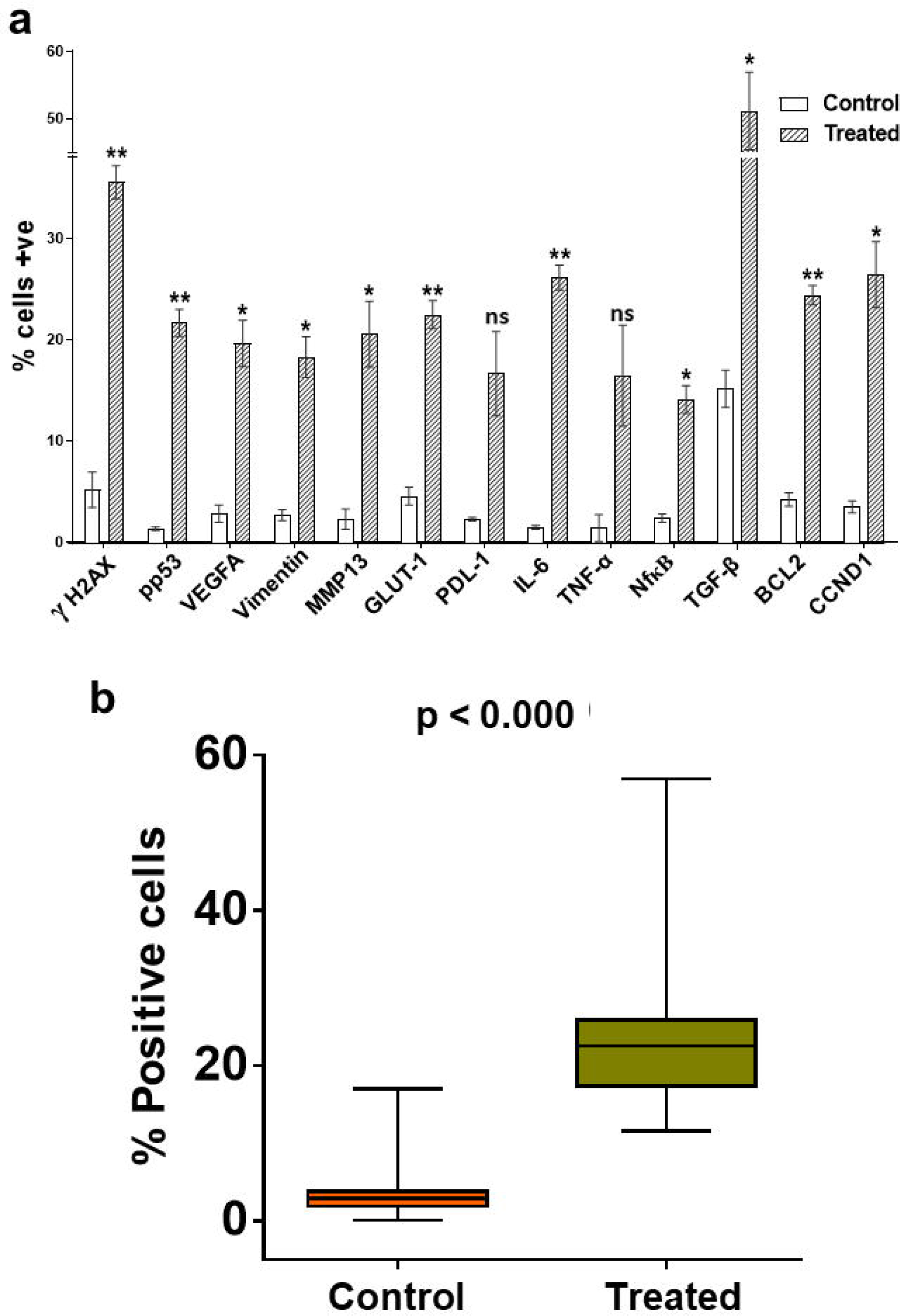
Activation of hallmarks of cancer in lung cells of SCID mice following intravenous injection of unlabelled MDA-MB-231 cells at 72 h. One hundred thousand unlabelled MDA-MB-231 cells were injected intravenously into SCID mice in the experimental group and saline was injected in the control group. Animals were sacrificed at 72 h and cryosections of the lungs were analysed by IF for 13 biomarkers representing 10 hallmarks of cancer; **a.** Histograms representing mean values (± SEM) of the percentage of cells positive for cancer biomarkers in MDA-MB-231-injected and control mice (n=2 in each group). For each biomarker, one thousand cells were analysed and the percentage of positive cells was calculated. The mean (± SEM) values in the control and MDA-MB-231-injected groups were compared using two tailed Student’s t test. **b.** Boxplots depicting comparative results of all biomarkers combined together. The median values were compared using the Mann–Whitney U test.

## Results

### Detection of fluorescently dually labelled cfChPs in the nuclei of lung cells

Fluorescently dually-labelled MDA-MB-231 and A-375 cells were injected intravenously into SCID mice and cryosections of their lungs were examined under fluorescent microscopy at 48 h. Representative images of fluorescently dual-labelled MDA-MB-231 and A-375 cells that were injected intravenously are given in Fig. 1a. Low power (10X) IF images of cryosections of lungs of the injected mice are given in Fig. 1b. The latter reveal patchy accumulation of co-localising red and green fluorescent signals representing cfChPs in the nuclei of the lung cells. The patchy nature of the signals is a reflection of the anatomy of the lung vasculature. The corresponding high power images (60X) are given in Supplementary Fig. 5 which again reveal the presence of dually-labelled cfChPs within the nuclei of lung cells. The presence of co-localising red and green fluorescent particles, rather than fluorescent intact cells, suggested that injected dually-labelled MDA-MB-231 and A-375 cells had died upon reaching the lungs and had released cfChPs which accumulated in the nuclei of lung cells.

### Intravenously injected cancer cells upregulate the hallmarks of cancer in lung cells

Since we had earlier observed that intravenous injection of B16F10 mouse melanoma cells leads to activation of two critical hallmarks of cancer viz. genome instability and inflammation in vital organs of mice when examined at 72 h **(9)**, we were curious to find out whether i.v. injection of MDA-MB-231 cells would activate cancer hallmarks in lung cells. IF analysis of cryosections of lungs of mice injected with (unlabelled) MDA-MB-231 cells at 72 h showed marked activation of all 13 biomarkers representing 10 hallmarks of cancer defined by Hanahan and Weinberg **(16)** (Figure 2a and Supplementary Figure 6). When compared statistically, 11 out of 13 biomarkers were found to be statistically significant, the exceptions being PDL-1 and TNF-α. A combined statistical analysis of all 13 hallmark biomarkers in the two groups revealed that the median value in the MDA-MB-231 injected groups was nearly eight-fold higher than that in the saline-injected controls (p < 0.000, Figure 2b). This finding indicated that the nuclear accumulation of cfChPs rapidly transformed mouse lung cells into incipient cancer cells by 72 h.

### Lung metastases contain both mouse- and human-specific DNA

Metastases detected on day 112 in case of MDA-MB-231 injected mice and on day 78 in case of A-375 injected mice were used for FISH analysis. FFPE sections were analysed by dual FISH by simultaneously using both mouse- and human-specific whole-genomic probes. Low power (10X) fluorescence images are given in Figure 3a in which almost the entire tumour area is clearly visible. The tumours are seen to contain both mouse DNA (green fluorescent signals) and human DNA (red fluorescent signals) in nearly equal quantities.

**Figure 3:**
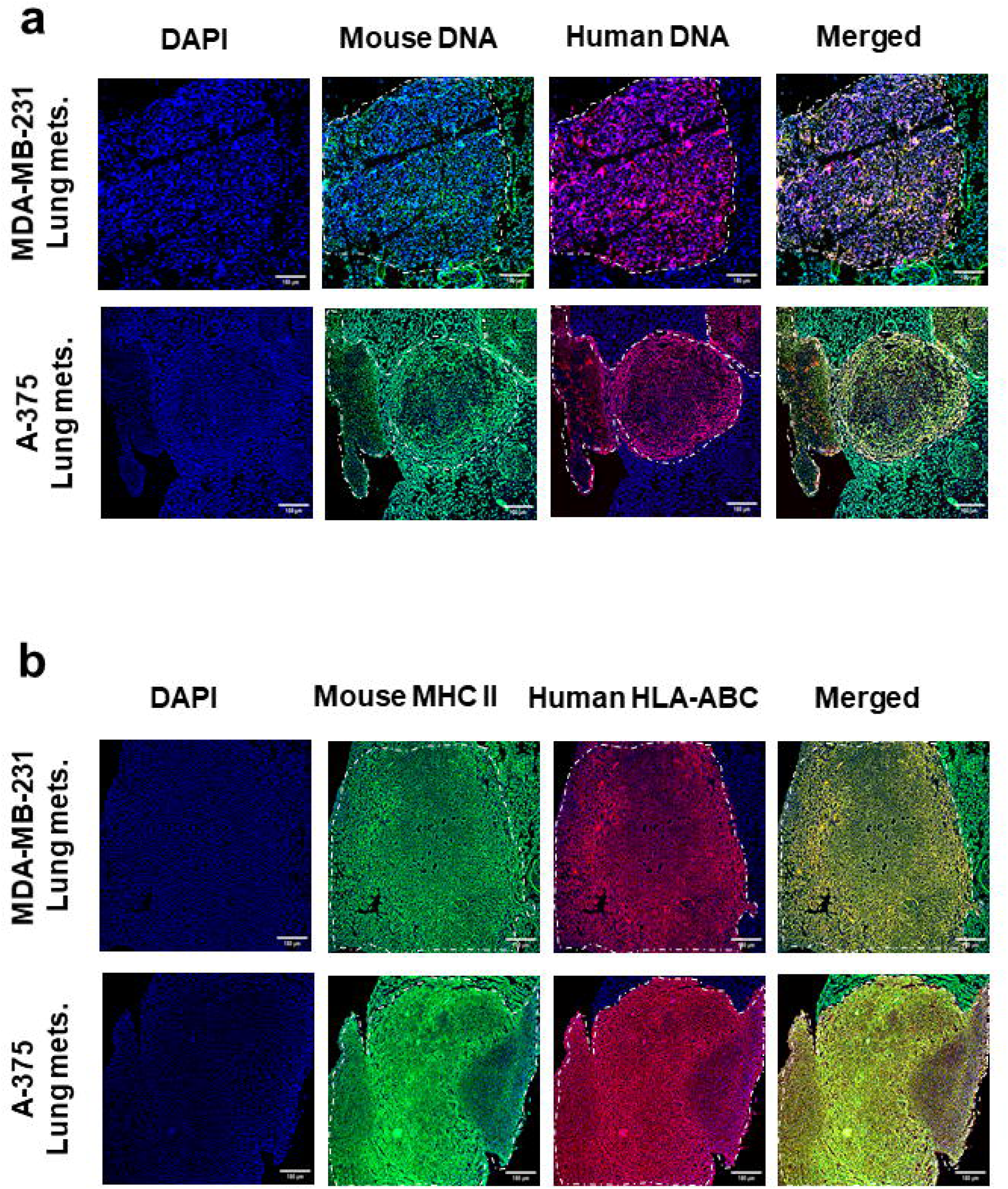
Detection of mouse and human specific DNA and of mouse and human specific proteins on FFPE sections of lung metastases in MDA-MB-231-and A-375-injected mice. **a.** FISH images following simultaneous use of mouse- and human-specific whole genomic probes. Presence of both mouse and human specific DNA are seen in the tumour cells. **b.** IF analysis using mouse and human specific mAbs (MHC-II and HLA-ABC, respectively). Presence of both mouse and human specific proteins are seen in the tumour cells.

The corresponding high power (60X) FISH images given in Supplementary Figure 7a confirm that the lung cell nuclei contain both mouse and human DNA. In order to quantify the FISH positive nuclei, one thousand DAPI stained nuclei were analysed for both MDA-MB-231 and A-375 induced tumours, and the percentage of FISH signals corresponding to mouse alone (green), human alone (red) and, mouse and human double-positive (yellow) signals were determined. The results obtained are as follows. For MDA-MB-231 induced tumours: mouse alone (3.075%), human alone (2.282%) and mouse and human double positives (94.643%). For A-375 induced tumours: mouse alone (1.101%), human alone (1.835%) and mouse and human double positives (96.881%).

### Lung metastases express both mouse- and human-specific proteins

The FFPE tumour sections were simultaneously immune-stained with monoclonal antibodies against species-specific mouse and human proteins (MHC-II and HLA-ABC, respectively). Low power (10X) IF images are given in Figure 3b in which almost the entire tumour area is clearly visible. The tumours are seen to express both mouse MHC-II (green fluorescent signals) and human HLA-ABC (red fluorescent signals) in nearly equal quantities.

The corresponding high power (60X) IF images are given in Supplementary Figure 7b which confirms that the lung cells were co-expressing mouse and human proteins. In order to quantify the mouse MHC-II, and human HLA-ABC expressing cells, one thousand DAPI stained cells were analysed for both MDA-MB-231 and A-375 induced tumours, and the percentage of protein expressing cells corresponding to mouse alone (MHC-II, green), human alone (HLA-ABC, red) and, mouse and human co-expressing (yellow) cells were determined. The results obtained are as follows. For MDA-MB-231 induced tumours: mouse alone (3.120%), human alone (2.340%) and mouse and human co-expressing cells (94.540%). For A-375 induced tumours: mouse alone (4.854%), human alone (4.660%) and mouse and human co-expressing cells (90.485%).

### Metastatic tumour cells in culture contain chimeric chromosomes

Metaphase spreads prepared from primary cultures of MDA-MB-231- and A-375-induced lung metastases, and their respective single-cell clones (MG6 and AG5), were examined by dual FISH i.e. by the simultaneous use of mouse and human-specific whole genomic probes. Both the primary cultures and the MG6 and AG5 clones revealed chimeric chromosomes with the chromosomal arms containing both human and mouse DNA (Figure 4a and 4b). The presence of human DNA in mouse chromosomes indicated that human cfChPs derived from dying MDA-MB-231 and A-375 cells had integrated and fused with mouse lung cell chromosomes.

**Figure 4:**
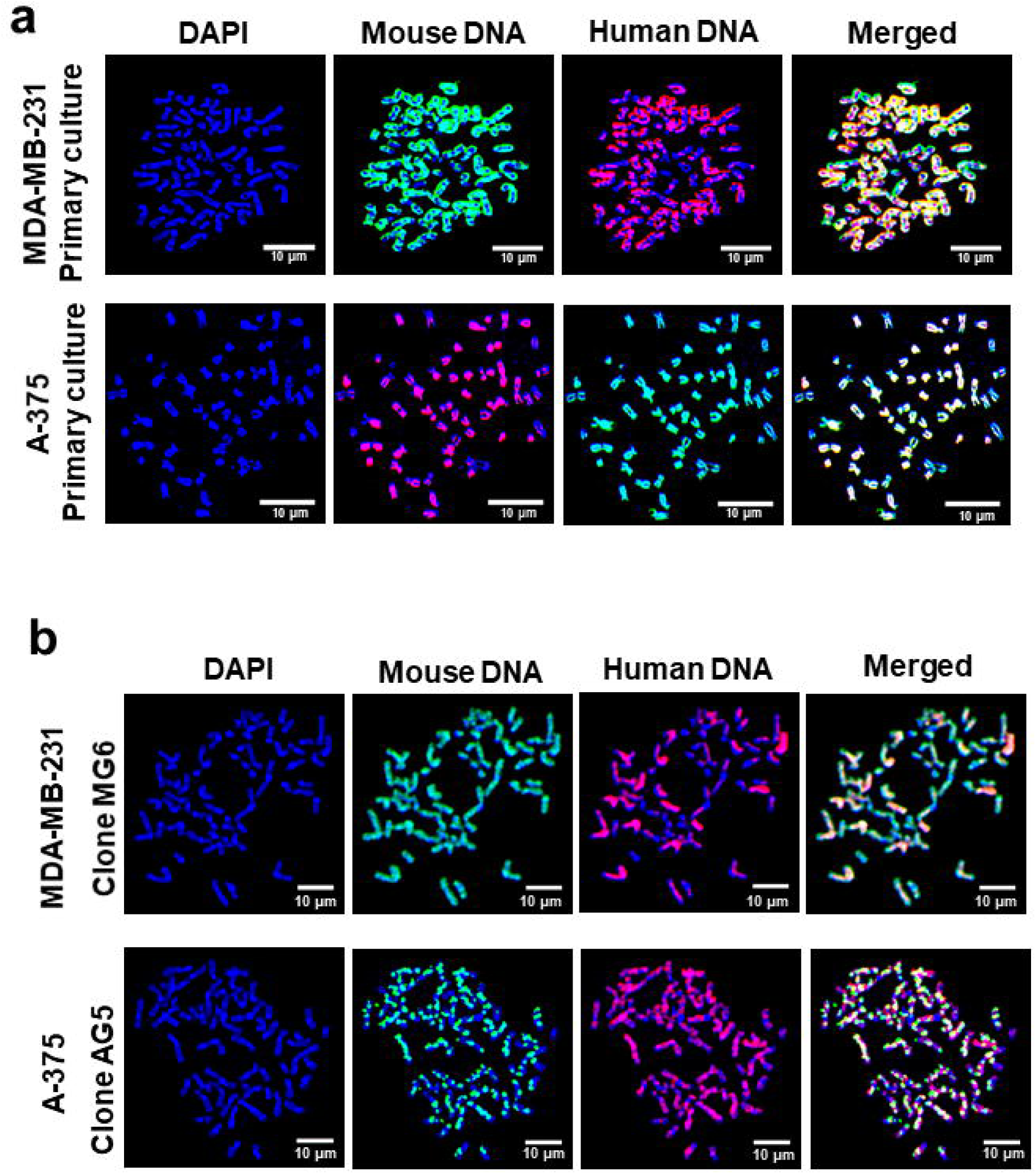
Detection of chimeric chromosomes in metaphase spreads prepared from cultured cells of MDA-MB-231 and A-375 lung metastases. **a.** Metaphase spreads prepared from cells of primary cultures analysed by FISH using human- and mouse-specific DNA probes. **b.** Metaphase spreads prepared from single-cell clones analysed by FISH using human- and mouse-specific DNA probes.

### Metastatic tumour cells in culture express both human and mouse proteins

Primary cultures of MDA-MB-231 and A-375 induced metastatic tumours, and MG6 and AG5 clones, were simultaneously immune-stained with monoclonal antibodies against MHC-II and HLA-ABC. Fluorescence microscopy revealed that the same cultured tumour cells expressed both human- and mouse-specific proteins (Figure 5a and 5b). This indicated that the human HLA-ABC gene that was carried via cfChPs to integrate into the mouse lung cell genomes was aberrantly expressing the human-specific HLA-ABC protein in the metastatic tumour cells that had developed in the mouse lung.

**Figure 5:**
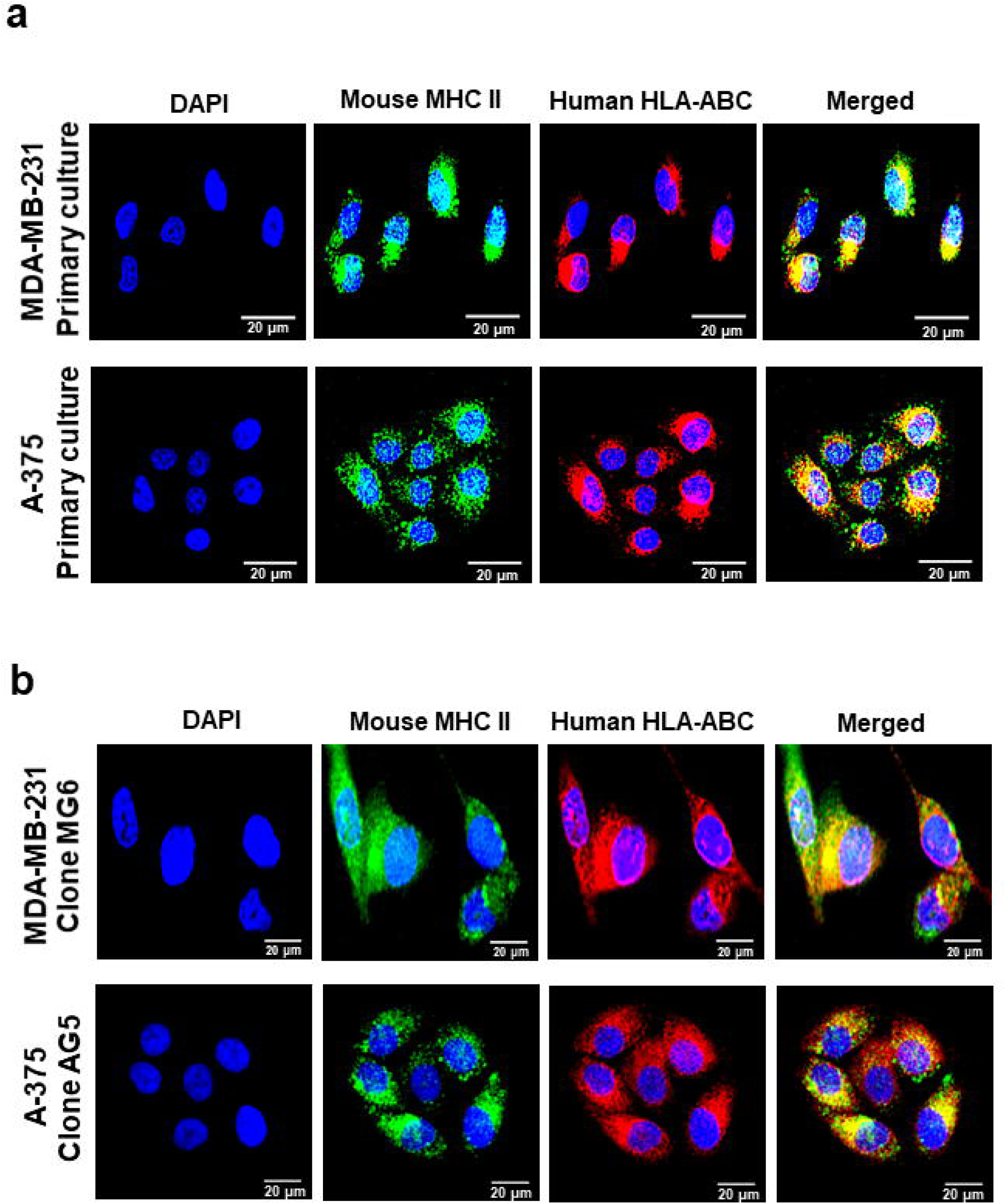
Co-expression of mouse- and human-specific proteins by the same cells of cultured MDA-MB-231 and A-375 lung metastases. **a.** Cells of primary cultures immune-stained with mouse- and human-specific mAbs (MHC-II and HLA-ABC, respectively). **b.** Clones MG6 and AG5 immune-stained with mouse- and human-specific mAbs (MHC-II and HLA-ABC, respectively).

**Figure 6:**
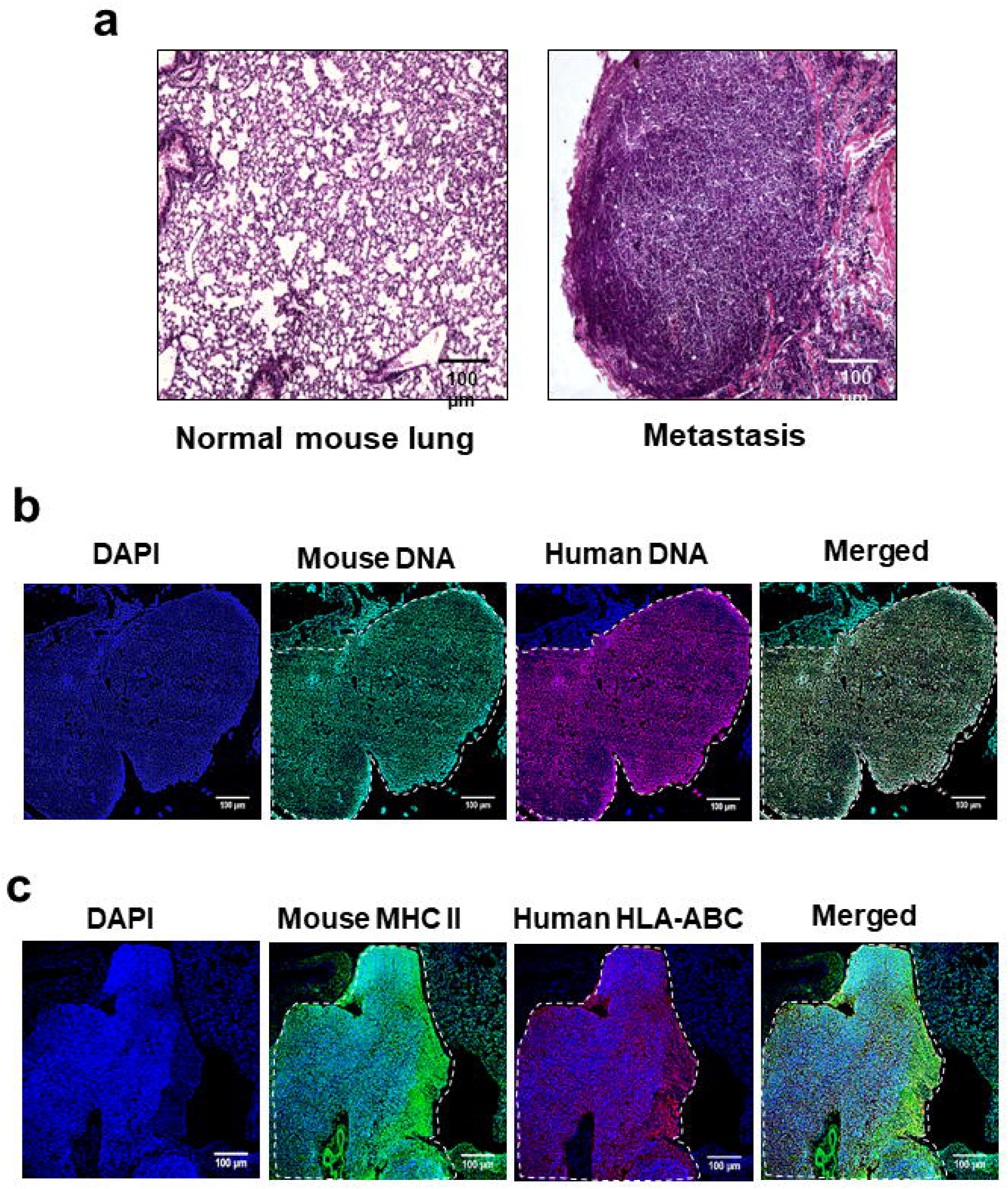
Development of lung metastases following i.v. injection of purified cfChPs isolated from radiation killed MDA-MB-231 cells and detection of mouse- and human-specific DNA and proteins in FFPE sections of the tumour. **a.** H&E sections showing lung metastases following i.v. injection of 700 ng of purified cfChPs. **b.** Detection of mouse- and human-specific DNA in lung metastasis. **c.** Detection of mouse- and human-specific proteins in lung metastasis.

### cfChPs isolated from radiation-killed MDA-MB-231 cells induce lung metastasis

As mentioned in the Methods section, two of five mice injected with cfChPs purified from radiation-killed MDA-MB-231 cells developed lung metastases on day 150. A H&E image of FFPE section of such a tumour is given in Figure 6a. The tumour sections were analysed with the simultaneous use of mouse and human specific whole genomic probes. A low power (10X) image given in Figure 6b shows DNA signals specific to both species to be present in the tumour with a preponderance of mouse DNA.

The corresponding high power (60X) fluorescent image shows vast preponderance of mouse DNA suggesting that human DNA contained in the purified cfChPs had integrated in the mouse lung cells and were involved in transforming them to generate a metastatic tumour (Supplementary Figure 8a). In order to quantify the FISH positive nuclei, one thousand DAPI stained nuclei were analysed, and the percentage of FISH signals corresponding to mouse alone (green), human alone (red) and, mouse and human double-positive (yellow) signals were determined. The results obtained are as follows: mouse alone (1.576%), human alone (0%) and mouse and human double positives (98.424%).

The cfChPs induced tumours were next analysed by IF with the simultaneous use of MHC-II and HLA-ABC mAbs. A low power (10X) image is provided in Figure 6c wherein the expression of both mouse- and human-specific signals are clearly seen with a preponderance of mouse proteins. A high power (60X) image is provided in Supplementary Figure 8b. In order to quantify the mouse MHC-II, and human HLA-ABC expressing cells, one thousand DAPI stained cells were analysed, and the percentage of protein expressing cells corresponding to mouse alone (MHC-II, green), human alone (HLA-ABC, red) and, mouse and human co-expressing (yellow) cells were determined. The results obtained are as follows: mouse alone (1.186%), human alone (0.791%) and mouse and human co-expressing cells (98.024%).

An unexpected and incidental finding was that of a tumourous outgrowths in the thymus region of three purified cfChP-injected mice (one out of five mice from the MDA-MB-231 group and both mice from the A-375 group; Supplementary Figure 9a). On H&E-stained sections, these tumours exhibited a highly undifferentiated pathology which precluded identification of their tissue origin (Supplementary Figure 9b). Nonetheless, FISH detected both human- and mouse-specific DNA in the same tumour cells, indicating that they had apparently been transformed by the genomic integration of cfChPs isolated from radiation-killed human cells (Supplementary Figure 9c). IF analysis also showed that the tumour cells co-expressed both HLA-ABC and MHC-II proteins (Supplementary Figure 9d).

## Discussion

Consistent with the findings of Fidler and others that the vast majority of tumour cells die following intravenous injection **(6,20,21)**, we observed that the injected MDA-MB-231 and A357 cells had also died upon reaching the lungs of the injected mice. We extensively analysed the cryosections of the lungs for presence of intact fluorescent cells in experiments in which fluorescently dually-labelled MDA-MB-231 and A-375 cells had been injected. An intact fluorescently dually-labelled cell should occupy an entire DAPI stained blue nucleus. Instead, what we detected were dually-labelled fluorescent particles representing cfChPs within the DAPI stained nuclei. This finding is consistent with our recent report wherein we had similarly injected fluorescently dually-labelled MDA-MB-231 cells into SCID mice and had looked for intact fluorescently dually-labelled cells in their brains. Here too, we had detected dually-labelled cfChPs within the nuclei of brain cells when examined at 72 h and not intact cells **(22)**. Nonetheless, even if we had missed detecting the presence of a few intact cells, the extensive presence of mouse DNA in the metastatic tumours strongly argues against the possibility that a few surviving intact human cancer cells that we had missed were responsible for metastasis formation. If the latter were to be true, the entire tumour should have comprised of human DNA. Our finding of chimeric chromosomes containing both mouse and human DNA suggests that cfChPs from dying human cancer cells had amalgamated with the genomes of mouse lung cells and were involved in the metastatic process. The detection of co-expression of both mouse MHC-II and human HLA-ABC by the same cultured cells further suggested that the genes contained within the chimeric chromosomes were being actively expressed. The most direct evidence that cfChPs were responsible for tumour formation comes from our demonstration that i.v. injection of purified cfChPs isolated from radiation-killed MDA-MB-231 cells also led to the formation of metastasis which predominantly contained mouse DNA. Taken together the above findings lead us to conclude that cfChPs released from dying cancer cells integrate into genomes of target cells to transform them to form new tumours which masquerade as metastasis. Such a conclusion is consistent with our hypothesis that metastasis arise from cells of target organs and not from those of the primary tumours **(9)**.

Does injection of large amounts of normal DNA promote carcinogenesis? To answer this question, it would have been desirable to undertake experiments wherein large amounts normal DNA, such as from normal mammary epithelial cells, were injected into SCID mice. This would have confirmed whether dying normal cells can also release cfChPs to integrate into genomes of mouse cells, and would have further substantiated the role of cfChPs in tumour metastasis. However, since our study was designed to investigate the mechanism by which the well-established models of MDA-MB-231 and A375 induced lung metastasis develop, experiments using normal cells were not conducted.

We detected marked activation of 13 biomarkers representing 10 hallmarks of cancer that have been defined by Hanahan and Weinberg **(16)** in lung cells of mice at 72 h following i.v. injection of MDA-MB-231 cells. This indicated that cfChPs had rapidly transformed the mouse lung cells into incipient cancer cells which ultimately went on to form detectable tumours after 2-3 months. These findings are consistent with our recent report that cfChPs are responsible for activation of cancer hallmarks and immune check-points in patients with advanced squamous cell carcinoma of the oral cavity **(23)**. We have further reported that cfChPs are directly responsible for activation of multiple immune check-points in isolated human lymphocytes **(24)**.

The new model of cancer metastasis suggested by our findings circumvents many of the problems associated with the conventional model in explaining the various steps of the metastatic cascade. For example, cfChPs are not susceptible to immune-surveillance or to the shearing force of circulation; they can freely negotiate the fine vascular capillaries, cross the blood-brain barrier and integrate into target cell nuclei.

The latter capabilities of cfChPs have been confirmed in our recent report cited above which clearly showed that cfChPs can readily cross the blood-brain-barrier and accumulate in the nuclei of brain cells **(22)**. The ability of the new model to circumvent the above mentioned obstacles makes it a preferred model of cancer metastasis than the conventional one.

Numerous studies have examined the relationship between primary and metastatic tumours at the genomic level. Overall, the results have been conflicting, with multiple studies reporting concordance **(25-28)** and others reporting discordance **(29-32)** between the profiles of primary and metastatic tumours. Significant similarities as well as differences in gene expression patterns have also been reported **(33-38)**. Studies investigating clonal evolution of metastases have also reported conflicting results. A wide repertoire of somatic changes in driver genes were reported which were either unique to the primary tumour or to the metastases or to be present in both **(39-46)**. Our finding that metastatic tumours contained DNA of both mouse and human origins and expressed both mouse and human proteins may help reconcile the above conflicting reports by suggesting that genetic signatures and gene expression patterns of both primary tumour cells and those of the target organs are to be expected in metastatic tumours. Conflicting reports in the literature may reflect the inherent bias of investigators while interpreting their results.

Our study could be criticized on the ground that we have depended largely on FISH and IF data, and that it would have benefited from genomic analyses to corroborate our findings. For example, DNA sequencing of the lung tumour cells and their respective single cell clones would have helped to illuminate what were the driver genes in the mouse lung metastases when MDA-MB-231 or A375 cells were injected intravenously? Comparison of DNA sequences of clones MG6 and AG5 with the injected MDA-MB-231 and A375 cells would have indicated the extent to which human DNA had integrated into the mouse lung tumour cells. Such investigations would have also elucidated on the conflicting reports in the literature alluded to above with respect to genomic and transcriptomic profiles of primary and metastatic tumours.

Nonetheless, the results of this study, in combination with our previous work in the field **(9,17,22-24)**, provide an alternative model to explain the fundamental pathophysiology of cancer metastasis, in which metastases arise *de novo* from the cells of the target organs and implicate cfChPs released from dying circulating tumour cells. The alternative model of metastasis is graphically illustrated in Figure 7.

**Figure 7:**
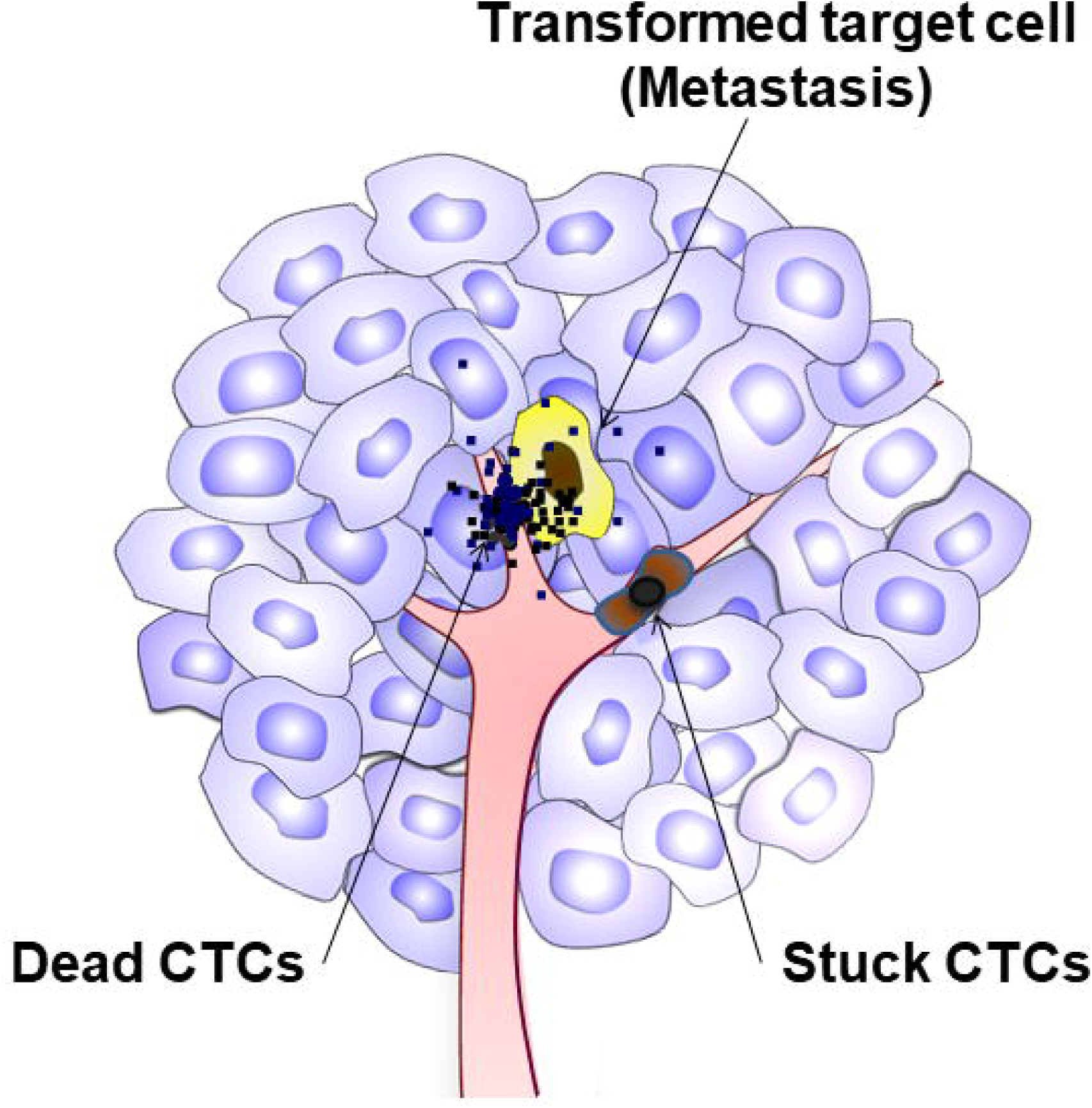
Graphical illustration of the alternative model of cancer metastasis. CTC = circulating tumour cells. The black dots represent cfChPs.

There is increasing recognition that therapeutic interventions can mobilise tumour cells into the circulation which could potentially contribute to metastatic dissemination **(47)**. This concern has been raised with respect to all three therapeutic modalities viz. chemotherapy **48,49)**, radiotherapy **50,51)** and surgery **52,53)**. We have recently reported that therapeutic interventions on human breast cancer xenografts promote systemic dissemination of oncogenes **(22)**. Xenografts of MDA-MB-231 cells generated in SCID mice were analyzed by IF and FISH to detect the presence of cfChPs in their brain cells. Multiple human DNA signals representing cfChPs were detected in nuclei of brain cells which co-localized with eight human onco-proteins. The number of co-localizing human DNA and human c-Myc fluorescent signals increased markedly after treatment with chemotherapy, localized radiotherapy or surgery. However, the signals could be dramatically minimized following concurrent treatment with three different cfChPs deactivating agents. Therefore, to prevent iatrogenic morbidity and mortality, it is imperative to re-examine the currently accepted mechanisms of cancer metastasis so that appropriate therapeutic approaches can be developed to prevent cancer dissemination. Before such a fundamental change occurs, however, further corroborative evidence in clinical settings is required to confirm our findings reported in the current study.

### Data availability

All data generated during this study are included in this published article.

### Ethics Statement

The experimental protocol was approved by the Institutional Animal Ethics Committee (IAEC) of the Advanced Centre for Treatment, Research, and Education in Cancer (ACTREC), Tata Memorial Centre (TMC), Navi Mumbai, India. The experiments were conducted in compliance with the ethical regulations and humane endpoint criteria of the IAEC and ARRIVE guidelines.

### Author contributions

GVR. designed and performed the experiments, analysed and interpreted the data, and wrote the manuscript. KP. supervised the experiments and analysed the data. SS, NKK, VJ, SS, RP, and HW. performed the experiments. IM. conceptualized and designed the project, was responsible for overall supervision and acquisition of funding, interpreted the data, wrote the manuscript, and approved the final draft.

### Funding

This study was supported by the Department of Atomic Energy, Government of India, through its grant CTCTMC to Tata Memorial Centre awarded to IM. The funding agency had no role in research design, collection, analysis, and interpretation of data and manuscript writing.

## Acknowledgements

We thank Mr. Ashish Pawar for his help in preparing this manuscript.

## Conflict of interests

The authors declare no competing interests.

## Notes

### Competing Interest Statement

The authors have declared no competing interest.

### Summary of Updates

1. Low power images (10X) of Figure 1b, Figure 3a & 3b and Figure 6b & 6c have now been provided. The earlier high power images (60X) have now been transferred as Supplementary Figure 5; Supplementary Figure 7a & 7b and Supplementary Figure 8a & 8b. 2. The earlier Supplementary Figure 5 has now been placed as Figure 6a. 3. In the new version we have provided quantitative data on percentage of fluorescent signals showing mouse alone, human alone, and mouse and human double-positive cells.

