## Supplementary material for "Evidence that metastases arise *de novo* as new cancers from cells of target organs": Legends to Supplementary Figures

**Supplementary Fig. 1:** Representative images of H&E sections of lungs showing presence of metastases in SCID mice injected with MDA-MB-231 and A-375 cells.

**Supplementary Fig. 2:** IF images showing epithelial nature of primary cultured cells and single-cell clones derived from MDA-MB-231 and A-375 lung metastases. **a&b.** MDA-MB-231 and A-375 primary cultured cells and the corresponding clones show positive epCAM staining. It is to be noted that, although A-375 is a human melanoma cell line, the primary cultures and clone AG5 derived from the A-375 induced lung metastasis show positive epCAM staining. This suggested that A-375 induced tumour cells were “epithelialized” by incorporating the epCAM gene from the mouse lung epithelial cells and were expressing the epCAM protein on their cell surfaces. This issue will become clear upon subsequent reading of the Results section of the manuscript.

**Supplementary Fig. 3:** Representative FISH and IF images showing lack of cross-reactivity between human and mouse specific DNA probes, and human and mouse specific mAbs. **a.** FISH images following simultaneous use of human (red) and mouse (green) specific DNA probes; **b.** IF images following simultaneous use of human (red) and mouse (green) specific mAbs.

**Supplementary Fig. 4:** Representative EM image of cfChPs purified from a culture medium of radiation (10 Gy) killed MDA-MB-231 cells. Beads-on-a-string appearance that is typical of chromatin is clearly seen. Histone octamers measuring ~50 nm is indicated by scale bar.

**Supplementary Fig. 5:** High power (60X) fluorescence microscopy images showing copious accumulation of fluorescently dually-labelled particles representing cfChPs within the nuclei of lung cells following intravenous injections of dual-labelled MDA-MB-231 and A375 cells at 48 h. One hundred thousand cells were injected intravenously into SCID mice in both the cases.

**Supplementary Fig. 6:** Representative IF images showing up-regulated cancer hallmarks in lung cells of SCID mice injected with MDA-MB-231 cells at 72 h.

**Supplementary Fig. 7:** High power (60X) fluorescent microscopy images of FFPE sections of lung metastases of MDA-MB-231- and A375-injected mice showing co-localising signals of mouse and human specific DNA and co-expression of mouse and human specific proteins. **a.** High power (60X) FISH images of lung metastasis simultaneously using mouse- and human-specific whole genomic probes. Co-localisation of mouse- and human-specific DNA is seen in the metastatic tumour cell nuclei. It should be noted that some of the FISH signals may not have been detected in the nuclei. This is because FISH analysis was done on FFPE sections, and the tissue cut surfaces may not always have been perfectly even leading to inconsistent binding of the FISH probes. **b.** High power (60X) IF images using mouse and human specific mAbs (MHC-II and HLA-ABC, respectively). Co-expression of MHC-II and HLA-ABC by the same cell is clearly visible in the metastatic tumour cells.

**Supplementary Fig. 8:** FISH and IF analysis of FFPE sections of lung metastases that developed following intravenous injection of purified cfChPs isolated from radiation-killed MDA-MB-231 cells showing co-localising signals of mouse- and human-specific DNA and co-expression of mouse- and human-specific proteins. **a.** Co-localising signals of mouse- and human-specific DNA; **b.** Co-expression of mouse- and human-specific proteins.

**Supplementary Fig. 9:** Image of a tumorous outgrowth in the thymus region showing the presence of both mouse- and human-specific DNA, and mouse- and human-specific proteins in tumour cells. **a.** Representative image of the tumourous growth; **b.** H&E image of the tumour; **c.** Dual FISH image on FFPE sections of the tumourous growth showing co-localisation of mouse- and human-specific DNA; **d.** Dual IF image on FFPE sections of the tumourous growth showing co-expression of mouse- and human-specific proteins.
