## Supplementary Table 1 for "Evidence that metastases arise *de novo* as new cancers from cells of target organs"

A. **FISH Probes**

| **Sr. No.** | **Reagent / FISH Probe** | **Catalogue No.** | **Company / Vendor** |
| --- | --- | --- | --- |
| 1. | Human Genomic DNA (probe) Red | Custom synthesized | Applied Spectral Imaging, Israel |
| 2. | Mouse Genomic DNA (probe) Red | Custom synthesized | Applied Spectral Imaging, Israel |

**B. Primary antibodies**

**Cancer hallmark antibodies**

| **Hallmark** | **Corresponding Marker** | **Catalogue No.** | **Company / Vendor** |
| --- | --- | --- | --- |
| **Sustained Proliferation** | TGF-β | T0438 | Merck-Millipore-Sigma, USA |
| **Evading Growth Suppressors** | pp53 | sc-18078 | Santa Cruz Biotechnology, USA |
| **Avoiding Immune Destruction** | PDL-1 | PRS-4059 | Merck-Millipore-Sigma, USA |
| **Enabling Replicative Immortality** | CCND-1 | sc-246 | Santa Cruz Biotechnology, USA |
| **Tumor Promoting Inflammation** | TNF-α | ab9739 | Abcam, UK |
|  | IL-6 | NB600-1131 | Novus Biologicals, USA |
|  | NFκB | ab32536 | Abcam,UK |
| **Invasion and Metastasis** | MMP13 | sc-12363 | Santa Cruz Biotechnology |
|  | Vimentin | #5741 | Cell Signaling Technology |
| **Inducing Angiogenesis** | VEGFA | ab 52917 | Abcam, UK |
| **Genome Instability and Mutation** | γH2AX | 05-636 | Merck-Millipore-Sigma, USA |
| **Resisting Cell Death** | BCL-2 | sc-492 | Santa Cruz Biotechnology,USA |
| **Deregulating Cellular Energetics** | Glut-1 | ab652 | Abcam, UK |

**Miscellaneous antibodies**

| **Sr. No.** | **Primary antibody** | **Catalogue No.** | **Company / Vendor** |
| --- | --- | --- | --- |
| 1. | BrdU | ab6326 | Abcam, UK |
| 2. | HLA -ABC class I | ab70328 | Abcam, UK |
| 3. | MHC class II | ab25333 | Abcam, UK |
| 4. | EpCAM | ab221552 | Abcam, UK |

**C. Secondary antibodies;**

| **Sr. No.** | **Secondary antibody** | **Company / Vendor** | **Catalogue No.** |
| --- | --- | --- | --- |
| 1. | Goat anti-rat IgG H&L (DyLight® 550) | Abcam, UK | ab98387 |
| 2. | Anti-mouse secondary antibody labeled with TRITC | Abcam UK | ab6785 |
| 3. | Anti-rat secondary antibody Fluorescein | Vector Labs, USA | FI-4000 |
| 4. | FITC-labeled anti-rabbit secondary antibody | Abcam®, UK. | ab6717 |
| 5. | FITC-labeled anti-mouse secondary antibody | Abcam®, UK. | ab6785 |
| 6. | FITC-labeled anti-goat secondary antibody | Abcam®, UK. | ab7121 |
| 7. | Donkey anti-goat Texas Red | Abcam® UK. | ab6883 |
| 8. | Rabbit anti-mouse IgG Rhodamine | Merck Millipore, USA. | SAB3701140 |
